# New Drosophila models to uncover the intrinsic and extrinsic factors mediating the toxicity of the human prion protein

**DOI:** 10.1101/2021.06.24.449794

**Authors:** Ryan R. Myers, Jonatan Sanchez-Garcia, Daniel C. Leving, Pedro Fernandez-Funez

## Abstract

Misfolded conformations of the prion protein (PrP) are responsible for devastating neurological disorders in humans and mammals. An unresolved problem is unraveling the mechanisms governing PrP conformational dynamics, misfolding, and the cellular mechanism leading to neurodegeneration. The variable susceptibility of mammals to prion diseases can be exploited to understand the conformational dynamics of PrP. Here we present a new fly model expressing human PrP with robust phenotypes in brain neurons and the eye. Using comparable attP2 insertions, we demonstrate the heightened toxicity of human PrP compared to that of mouse and hamster PrP along with a specific interaction with the amyloid-beta peptide. Using this new highly toxic new model, we started to uncover the intrinsic (sequence / structure) and extrinsic (interactions) factors regulating PrP toxicity. As extrinsic factors, we describe the importance of the PERK - ATF4 branch of the unfolded protein response as a key cellular mechanism mediating the toxicity of human PrP. For intrinsic factors, we introduced point mutations in human PrP (N159D, D167S, N174S) that were partially protective, revealing its high propensity to misfold into toxic conformations.

## Introduction

Prion diseases encompass a clinically heterogeneous class of brain disorders in humans with direct molecular and pathological correlates in several mammals (Mathiason, 2017, Zlotnik and Rennie, 1965). The main pathological features shared by prion diseases in humans and animals are spongiform degeneration of the brain and accumulation of misfolded, insoluble conformations of the prion protein (PrP) (Colby and Prusiner, 2011, Scheckel and Aguzzi, 2018). PrP is a glycoprotein anchored to the extracellular aspect of the membrane highly expressed in brain neurons but not essential for survival (Sigurdson et al., 2019, Steele et al., 2007, Bueler et al., 1992). Other than humans, only ruminants suffer endemic prion diseases: scrapie in sheep and goats, and chronic wasting disease in deer and moose. In addition to these endemic prion diseases, several mammals proved susceptible to laboratory transmission (chimpanzee, mouse, hamsters, bank vole) (Chandler, 1971, Zlotnik and Rennie, 1965, Chandler and Fisher, 1963, Zlotnik and Rennie, 1963), but not rabbits (Gibbs and Gajdusek, 1973, Barlow and Rennie, 1976). The spread of contaminated bone meal carrying sheep prions revealed the susceptibility to prion diseases of cattle, felines, and mustelids (ferret family) (Kirkwood and Cunningham, 1994, Sigurdson and Miller, 2003). Notably, a few domestic animals exposed to the same contaminated feed demonstrated resistance to zoonotic prion transmission: dogs, horses, rabbits, and pigs (Kirkwood and Cunningham, 1994). Overall, these observations uncover natural differences in the susceptibility to prion diseases, which can be exploited to further understand the rules governing PrP misfolding and disease. The most likely mechanism responsible for differing animal susceptibilities to prion diseases is the presence of natural variations in the PrP sequence that alter its conformational dynamics. That is, disease susceptibility is encoded by intrinsic differences in amino acid sequence that modulate conformational dynamics and disease, without a relevant impact of the cellular milieu (Vorberg et al., 2003, Vilette et al., 2001). This knowledge can be leveraged to unravel how sequence variation (genotype) impacts PrP toxicity (phenotype) (Myers et al., 2020).

Over the last few years, we and others created a collection of transgenic *Drosophila* models expressing PrP from susceptible and resistant animals: Syrian hamster, mouse, sheep, rabbit, dog, and horse (Sanchez- Garcia and Fernandez-Funez, 2018, Fernandez-Funez et al., 2010, Fernandez-Funez et al., 2009, Gavin et al., 2006, Thackray et al., 2012b). These studies support the idea that the intrinsic properties of each PrP are preserved when expressed in flies: WT hamster, mouse, and sheep PrP are toxic in flies whereas WT rabbit, horse and dog are not (Sanchez-Garcia and Fernandez-Funez, 2018, Thackray et al., 2012a, Thackray et al., 2012b, Fernandez-Funez et al., 2010). These differences in toxicity correlate with PrP conformational dynamics, with rabbit, horse and dog PrP showing low misfolding and aggregation (Fernandez-Funez et al., 2010, Khan et al., 2010). Additionally, *Drosophila* demonstrates high sensitivity to subtle changes in the PrP sequence based on the observation that WT hamster PrP is more toxic than WT mouse PrP (Fernandez-Funez et al., 2010), and dog and horse PrP carrying humanized point mutations become toxic (Sanchez-Garcia and Fernandez-Funez, 2018). Sensitive behavioral and morphological assays document the progressive toxicity of rodent and sheep PrP coupled with age-dependent changes in conformation and aggregation (Fernandez-Funez et al., 2009, Thackray et al., 2012b). These relevant assays are time-consuming, which dramatically narrows the utility of the fly PrP models. *Drosophila* is an ideal tool for cost-effective and efficient gene discovery when robust, easy to score, and sensitive assays are available. The *Drosophila* eye offers unique advantages for conducting genetic screens since subtle changes in the eye size can be easily determined (Fernandez-Funez et al., 2000). Unfortunately, existing PrP models not toxic in the fly eye (Fernandez-Funez et al., 2017), limiting their application.

To expand the utility of *Drosophila* to uncover the mechanisms mediating PrP toxicity, we examined the possibility that PrP from other animals could be more toxic. We hypothesized that human PrP was likely to be more toxic than PrP from other mammals with endemic or acquired prion diseases (bovine, sheep, deer) for the following reasons. 1) Human prion diseases, unlike other animals, present with sporadic, genetic, and infectious etiologies, arguing for high structural instability. 2) Human prion diseases are clinically highly heterogenous neurological disorders that manifest with cognitive, movement, or sleep perturbations. To the best of our knowledge, endemic prion diseases have single presentations in each host. 3) These differences can be attributed to naturally diverse prion strains with specific neurotropisms, supporting the high conformational dynamics of human PrP. 4) Inherited prion diseases in humans are caused by more than 50 missense mutations, some of which introduce subtle structural changes (e.g., V180I, V210I), suggesting that minor perturbations in the sequence of human PrP can dramatically alter its dynamics. To test this idea, we generated flies expressing human PrP. To limit the risk that these flies could accumulate the transmissible, protease-resistant scrapie PrP (PrP^Sc^) conformation, we conducted this work at a BSL3 facility. We showed recently that flies expressing human PrP-V129 exhibit a powerful new phenotype - small and glassy eyes - that supports the heightened toxicity of human PrP (Fernandez-Funez et al., 2017). However, we could not directly compare the toxicity of the flies expressing human PrP against existing models due to differences in construct design affecting the expression levels.

Here, we describe additional novel phenotypes in the brain and in a behavioral assay induced by *<u>random</u>* human PrP-V129 and -M129 insertions. We also describe a new suite of comparable, isogenic transgenic flies carrying human and rodent PrP: codon-optimized and inserted in the same molecularly defined attP landing site (Bischof et al., 2007). These new *<u>attP2-based</u>* PrP models elegantly demonstrate the heightened toxicity of human PrP compared to hamster and mouse PrP. As proof-of-concept for the utility of the new human PrP models, we identified intrinsic and extrinsic factors modulating its toxicity. It is well documented that misfolded PrP in the ER triggers the unfolded protein response (UPR) (Hetz et al., 2007, Hetz et al., 2003), a complex pathway with both protective and maladaptive consequences (Hetz, 2012, Moreno et al., 2012). We describe here that PERK and ATF4 loss-of-function are protective in combination with PrP, indicating that PERK is a major driver of PrP toxicity. Lastly, to gain a mechanistic understanding of the sequence-structure determinants of human PrP toxicity, we introduced three protective mutations from animals resistant to prion diseases (Sanchez-Garcia and Fernandez-Funez, 2018). D167S and N174S partially suppress human PrP toxicity whereas N159D does not, illustrating the high disease propensity of human PrP. These improved *Drosophila* models of proteinopathies provide expanded opportunities to identify the intrinsic and extrinsic factors mediating PrP toxicity, including high-throughput genetic screens and targeted amino acid replacements to determine the rules governing PrP toxicity.

## Results

### Structural differences between human and rodent PrP

It is unclear, at this time, which differences at the sequence and structural levels contribute to the natural risk of developing prion diseases and to the phenotypic differences observed in transgenic flies (Fernandez-Funez et al., 2010, Fernandez-Funez et al., 2009, Fernandez-Funez et al., 2017). The sequence alignment of the globular domain of human PrP demonstrates extensive similarity to that of hamster and mouse PrP with minor differences (Fig. 1A). All the sequences are numbered according to human PrP to avoid confusion. Most amino acid differences between human and rodent PrP are conservative (side chains with similar chemical properties). Interestingly, helix 2 and the first half of helix 3 are identical for the three sequences (Fig. 1A), whereas helix 1 displays only one amino acid difference. Most of the variation is concentrated in the loops and the end of helix 3. It is important to note that the highly variable region comprised of the loop between the *β*-sheet and helix 2 (*β*2-*α*2 loop) forms a 3D domain with the distal helix 3 (Fig. 1B). This domain is proposed to play a critical role in PrP misfolding and is the proposed binding site of a hypothetical protein that regulates PrP misfolding (Telling et al., 1995, Kaneko et al., 1997). For simplicity, we termed this region the C-terminal 3D (CT3D) domain (Fig. 1B). The 3D alignment of the globular domain of human and rodent PrP (Zahn et al., 2000, Calzolai et al., 2000, James et al., 1997, Riek et al., 1996) shows overt similarity (Fig. 1C and D). A close examination reveals mild differences that may underlie their distinct toxicity. For instance, human PrP has a longer *β*-sheet than rodent PrPs despite perfect sequence conservation, suggesting that the *β*-sheet is more stable in human PrP (Fig. 1D). Mouse PrP has a 310 turn in the *β*2-*α*2 loop that indicates its increased stability (Fig. 1C and D). Additionally, helix 2 starts at N173 in human PrP, Q172 in hamster PrP, and N171 in mouse PrP, resulting in a shorter helix in human PrP (Fig. 1C, arrow). Comparing the position of two conserved amino acids in the loop, mouse Y169 is deeply buried in the hydrophobic core, is highly exposed in human, and is in-between in hamster PrP (Fig. 1C). D167 is more exposed in human than in mouse and hamster, creating a more open loop. These subtle amino acid differences suggest that mouse is the most stable of these three structures and human is likely the most unstable, which informs our hypothesis.

**Figure 1.**
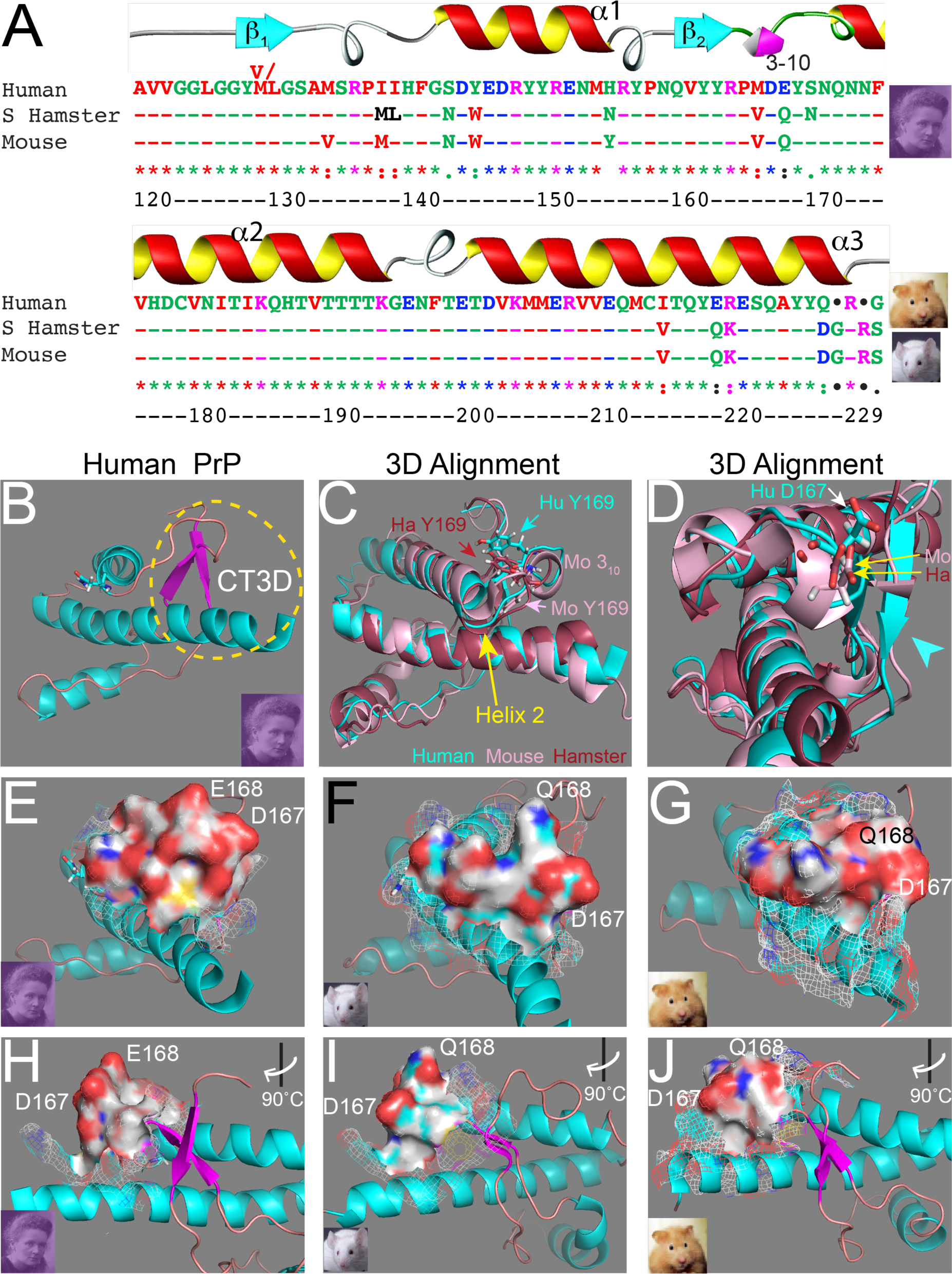
Sequence / structure differences between human and animal PrP. **A**, Sequence alignment of the C-terminal globular domain of PrP from human, Syrian hamster, and mouse. Amino acid numbering corresponds to human PrP throughout to avoid confusion. The alignment shows high overall conservation with most variation clustered in the *β*2-*α*2 loop and distal helix 3. **B**, 3D visualization of human PrP indicating the position of the CT3D domain (yellow dotted circle). **C and D**, 3D alignment of human (cyan), mouse (pink) and hamster (brown) globular domains. Overall, the three proteins share the same structural elements, with three helices and a short *β*-sheet. However, mouse has a 310 turn in the loop and helix 2 is longer (C, yellow arrow). The different position of Y169 in human, mouse, and hamster is indicated (C). The *β*-sheet is longer in human, followed by hamster and is shorter in mouse (D, arrowhead). The position of D167 is indicated for human, mouse, and hamster (D). **E-J**, Surface and Mesh views for the *β*2-*α*2 loop, front view (E-G) and side view (H-J). In human PrP, the loop is vertical and tall, with two acidic residues sticking upwards (E and H). In mouse PrP, the loop is not as tall, with D167 shifted to a lower position (F and I). In hamster PrP, the loop is flat and closer to helix 3 (G and J).

Surface visualization of the CT3D domain indicates that in human PrP, the side chains of D167 and E168 are perpendicular to helix 3, resulting in a positive charge (Fig. 1E and H). Most animals carry D167- Q168 in the equivalent positions (Fig. 1A), resulting in a less charged domain (Fig. 1F and G). In mouse PrP, Q168 is upward, but the rest of the loop is lower (Fig. 1F and I). Interestingly, the loop in hamster PrP is lower and flatter than in human and mouse PrP, resulting in a closer interaction with helix 3 (Fig. 1G and J). This visual analysis of 3D structures identifies significant differences in the CT3D domain supporting a more the increased conformational stability of rodent PrP compared to human PrP.

### New Drosophila eye phenotype of <u>random</u> human PrP insertions

We showed before that a random insertion of codon-optimized human PrP-V129 induces a new eye phenotype (Fernandez-Funez et al., 2017). We compare here the expression of codon-optimized human PrP-V129 and M129 from random insertions. M/V129 is a polymorphism of human PrP that is significant for the risk of variant Creutzfeldt-Jacob disease (CJD) transmission from cattle, but otherwise has no impact on the causation of other prion diseases (Kobayashi et al., 2015). Expression of PrP-V129 and M129 under the control of *GMR-Gal4* resulted in disorganized, glassy eyes (Fig. 2A-C), with PrP-M129 causing a smaller eye (Fig. 2C). To examine these flies in more detail, we fixed the eyes of 1-day-old flies, embedded them in resin and sectioned them. Semithin sections (1 μm thick) show that control flies expressing CD8- GFP at 27°C display a regular arrangement of ommatidia, the visual units of the compound eye (Fig. 2D). Most ommatidia contain seven photoreceptors, recognized for the specialized photosensitive rhabdomeres in the center that stain darker, although the high temperature and the intrinsic toxicity of Gal4 cause some disruption. Flies expressing human PrP-V129 have disorganized and vacuolated retinas (Fig. 2E). Most ommatidia contain fewer photoreceptors and their arrangement appears disrupted. Flies expressing human PrP-M129 show retinas with more prominent disorganization and vacuolation, and few recognizable rhabdomeres (Fig. 2F). To examine these phenotypes at cellular resolution, we collected ultrathin sections (70 nm thick), stained them and imaged them by transmission electron microscopy. Control flies show the normal polygonal arrangement of seven photoreceptors (R1-R7) around the rhabdomeres (Fig. 2G). These eyes contain fewer pigment granules in the surrounding cells but display relatively healthy ommatidia. Flies expressing PrP-V129 show rhabdomere loss and the remaining rhabdomeres are small and disorganized (Fig. 2H). One of the photoreceptors (*) appears vacuolated and others contain hyperplastic endoplasmic reticulum (ER) (Fig. 2H, arrowheads). Flies expressing PrP-M129 show few rhabdomeres and extensive vacuolation of photoreceptors (Fig. 2I, *). The rhabdomeres show low electron density and fusions, indicating severe perturbations. Lastly, most mitochondria appear vacuolated with disrupted internal membranes (Fig. 2I, m). Overall, the examination of the eyes in flies expressing random insertions for human PrP-V129 and M129 show strong perturbations of rhabdomere differentiation and cell survival.

**Figure 2.**
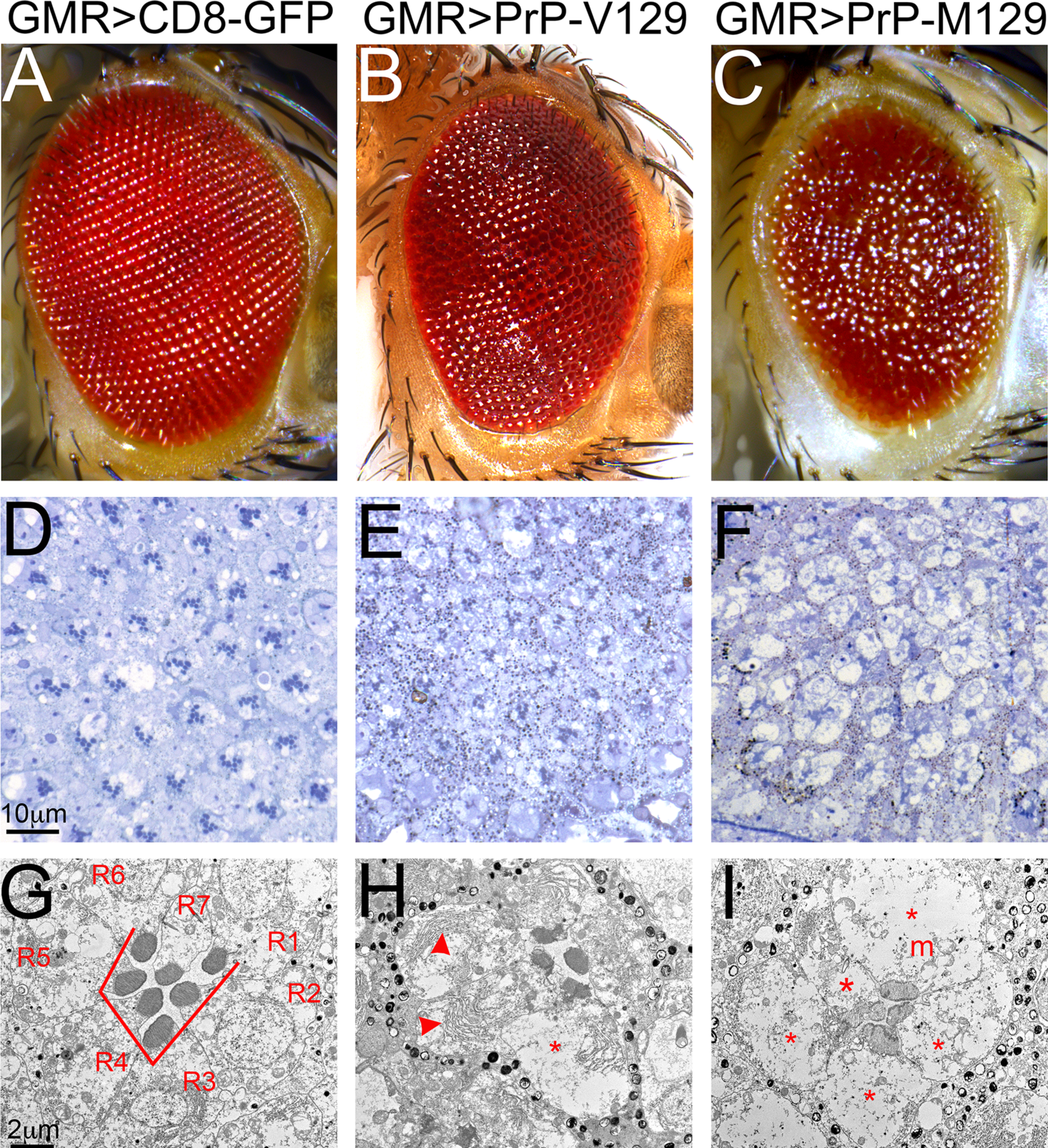
New eye phenotypes of random human PrP lines. **A-C**, Micrographs of fresh eyes expressing mCD8-GFP (A), human PrP-V129 (B), or human PrP-M129 (C) in the eye from random insertions under the control of *GMR-Gal4* at 27°C. Control flies and flies expressing hamster PrP exhibit highly organized eyes (A). Flies expressing human PrP-V129 or M129 display disorganized, and glassy eyes, but M129 seems to induce the stronger perturbations. **D-F,** Semithin sections of the retina. D, Expression of mCD8- GFP preserves the lattice of ommatidia and most ommatidia contain seven photoreceptors. E, Expression of PrP-V129 results in disorganized ommatidia and loss of photoreceptors as indicated by fewer than seven rhabdomeres per ommatidia. F, Expression of PrP-M129 results in disorganized and vacuolated retina with loss of photoreceptors. **G-I**, Transmission electron micrographs of single ommatidia. G, mCD8-GFP have seven photoreceptors (R1-R7) and preserves the organization of the rhabdomeres. H, Expression of PrP- V129 results in partial vacuolation of photoreceptors (*), abnormal rhabdomeres, and excess of ER (arrowheads). I, Expression of PrP-M129 results in highly vacuolated photoreceptors (*) and mitochondria (m), and hypochromic rhabdomeres.

### New brain phenotypes caused by <u>random</u> human PrP insertions

We next examine the consequence of expressing human PrP in brain neurons to identify new phenotypes. Flies constitutively expressing human PrP under the control of the pan-neural driver *Elav-Gal4* show 100% lethality at 25°C, meaning that no adult flies eclose. In contrast, flies expressing hamster PrP under the same conditions are 100% viable. To bypass this developmental toxicity, we used the *Elav-GeneSwitch* driver (*Elav-GS*), a conditional Gal4 activated by the steroid hormone mifepristone (RU486) (Roman et al., 2001). We combined LacZ (negative control), hamster PrP, and human PrP with *Elav-GS*, and grew the flies in media lacking RU486 to allow development in the absence of PrP expression. Then, we collected newly eclosed adult flies, placed them in vials with or without RU486 at 28°C (Day 0), and subjected the flies to climbing assays. Control experiments (- RU486) showed similar climbing ability in flies carrying LacZ, hamster PrP, or human PrP constructs (Fig. 3A). Flies expressing LacZ (+RU486) reached 50% climbing index by day 16 and climbed until day 28 (Fig. 3A). Flies expressing hamster PrP (+RU486) reached 50% climbing index at day 14 and continued to climb until day 26 (Fig. 3A). However, flies expressing human PrP (+RU486) reached 50% climbing index by day 1.5 and only climbed for 3 days (Fig. 3A). The fast progression of the locomotor dysfunction illustrates the hihg toxicity of human PrP.

**Figure 3.**
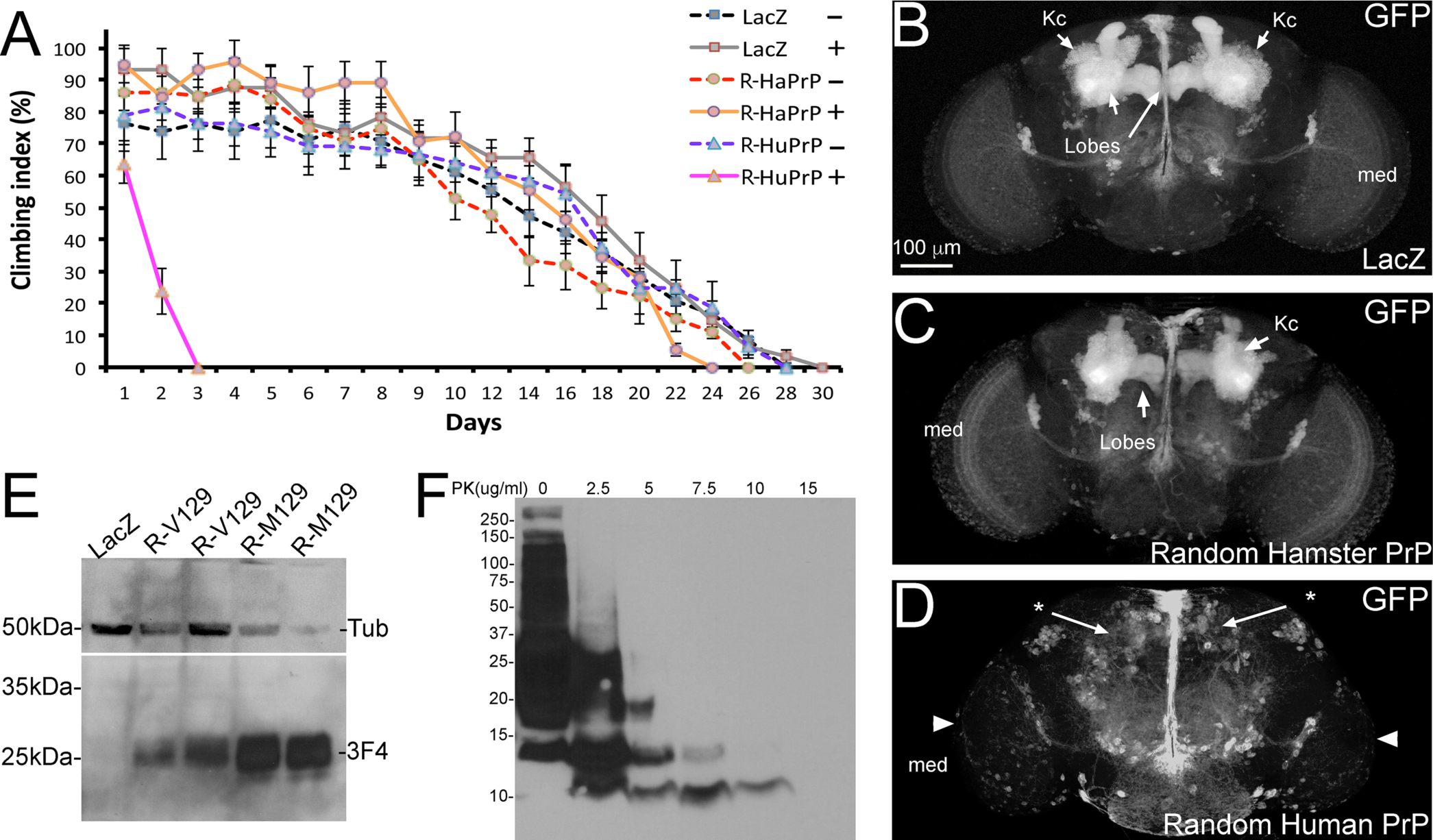
New phenotypes induced by human PrP in *Drosophila*. **A,** A <u>random insertion</u> for human PrP induces aggressive locomotor dysfunction. LacZ (squares), hamster PrP (HaPrP, circles), and human PrP (HuPrP, triangles) were conditionally expressed pan-neurally using *Elav-GS*. Expression was activated at day 1 for flies (+, continuous line) or not activated at all (-, broken line). Only flies expressing human PrP exhibit aggressive locomotor dysfunction. **B-D,** Whole brains showing the consequence of expressing human PrP in the mushroom bodies (*OK107-Gal4 / UAS-mCD8-GFP*). B, Expression of LacZ reveals large mushroom body (MB) clusters in 1-day-old flies, including large Kenyon cell clusters (Kc) and axonal projections. C, Expression of hamster PrP has no effect on the development of the MB complexes. D, Expression of human PrP-V129 completely eliminates the mushroom body clusters, leaving no traces of MB neurons and resulting in smaller optic lobes. Arrowhead points to the smaller medulla (med). **E,** Fly homogenates expressing LacZ (lane 1), human PrP-V129 (lanes 2 and 3), or human PrP-M129 (lanes 4 and 5) in the eye from random insertions under the control of *GMR-Gal4* at 27°C. The membrane was incubated with anti-Tubulin and 3F4 anti-PrP. V129 and M129 display similar electrophoretic mobility, but M129 accumulates at higher levels. **F,** Homogenates from 10-day-old fly heads expressing human PrP in the eye subjected to a mild PK gradient (30 min, 25°C). The 10 μg/ml treatment degraded almost all PrP except for a small fragment at 10 kDa. The 15 μm/ml digestion has no detectable PrP.

We next monitored the impact of constitutive expression of human PrP on the architecture of a brain center not critical for survival. The mushroom bodies are a well-known brain region involved in higher neural processing in insects, including memory and learning (Davis, 2005, Tanaka et al., 2008). The mushroom bodies are two symmetric centers consisting of around 2,500 neurons each with the cell bodies in the posterior brain and the axonal projections extending the front. We constitutively expressed LacZ or hamster PrP in mushroom body neurons under the control of *OK107-Gal4* and examined the architecture of the mushroom bodies in 1-day-old flies. These flies show robust mushroom bodies (Fig. 3B and C), although the axonal projections are slightly smaller in flies expressing hamster PrP (Fig. 3C). Notably, flies expressing human PrP have lost the mushroom bodies (Fig. 3D). The optic lobes are smaller due to weal expression of *OK107-Gal4* in the eye and medulla (Fig. 3D, arrowheads). Overall, these new phenotypes induced by human PrP in the fly brain and support our hypothesis that human PrP is more toxic than rodent PrPs. These phenotypes though are not directly comparable since only human PrP was codon-optimized for *Drosophila* and all these constructs are inserted in different loci.

### Protein analysis of <u>random</u> insertions of human PrP

We subjected homogenates from heads of 1-day old flies expressing LacZ (negative control), human PrP-V129, or -M129 under the control of *GMR*-*Gal4* to western blot and detected PrP with the 3F4 antibody. Duplicates for M129 and V129 show that M129 levels are several folds higher than that for V129, possibly explaining the difference in eye phenotype (Fig. 3E). This exemplifies the problem with random insertions. We next determined whether human PrP accumulates in protease resistant PrP conformations in *Drosophila*. Transmissible prions contain PrP^Sc^, which has unique properties, including high resistance to denaturing agents (urea, guanidinium) and to harsh proteinase-K (PK) digestion (20 μg/ml PK for 1h at 37°C). PK digestion of PrP^Sc^ results in a diagnostic PK-resistant core fragment of around 20 kDa that is transmissible. We expressed human PrP in the eye and aged the flies for 10 days. Then, we homogenized the heads, split the homogenate in aliquots, and subjected them to a mild PK gradient (2.5-15 μg/ml PK for 30 min at 25°C) to detect the presence of PrP^Sc^ (Fig. 3F). Incubation with 5 μg/ml PK eliminated all the full-length PrP but left several fragments below 20 kDa. Incubation with 7.5 and 10 μg/ml PK eliminated almost all the signal, except for small fragments around 12 and 10 kDa. Finally, incubation with 15 μg/ml PK eliminated all PrP signal. Thus, digestion under mild PK conditions demonstrate that human PrP shows relative resistance to proteolysis but no spontaneous formation of PrP^Sc^ in *Drosophila*.

### New human and rodent PrP constructs: codon-optimized <u>attP2</u> lines

To determine whether the structural differences between human and rodent PrP described above are responsible for the different toxicity *in vivo*, we generated a new, comparable suite of human and rodent PrP constructs. All the PrP constructs were *(a)* codon-optimized for *Drosophila* expression and *(b)* inserted in the same molecularly defined locus, the strong attP2 landing site that we used before (Bischof et al., 2007, Moore et al., 2018). For human PrP, we generated the two natural polymorphisms (M129 and V129) to examine their behavior when expressed from comparable insertions. These new constructs enable comparative studies in which any differences in toxicity between PrP constructs can be directly attributed to sequence differences. We expressed the new attP2 PrP constructs in the eye under the control of *GMR- Gal4*. Control flies expressing mCD8-GFP-attP2 exhibit highly organized eyes (Fig. 4A and F) with a regular ommatidia lattice (Fig. 4F, inset). Flies expressing mouse or hamster PrP-attP2 have normal eyes similar to those of control flies (Fig. 4B, C, G and H). Flies expressing human PrP-M129-attP2 or V129- attP2 show mild disorganization of the eye (Fig. 4D, E, I and J). High magnification shows poor organization and differentiation of ommatidia with multiple fusions (Fig. 4I and J, arrowheads). The eye phenotypes of the two human PrP-attP2 lines are weaker than those from random insertions (Fig. 2C), which is expected when comparing strong random insertions identifies lines with the strong attP2.

**Figure 4.**
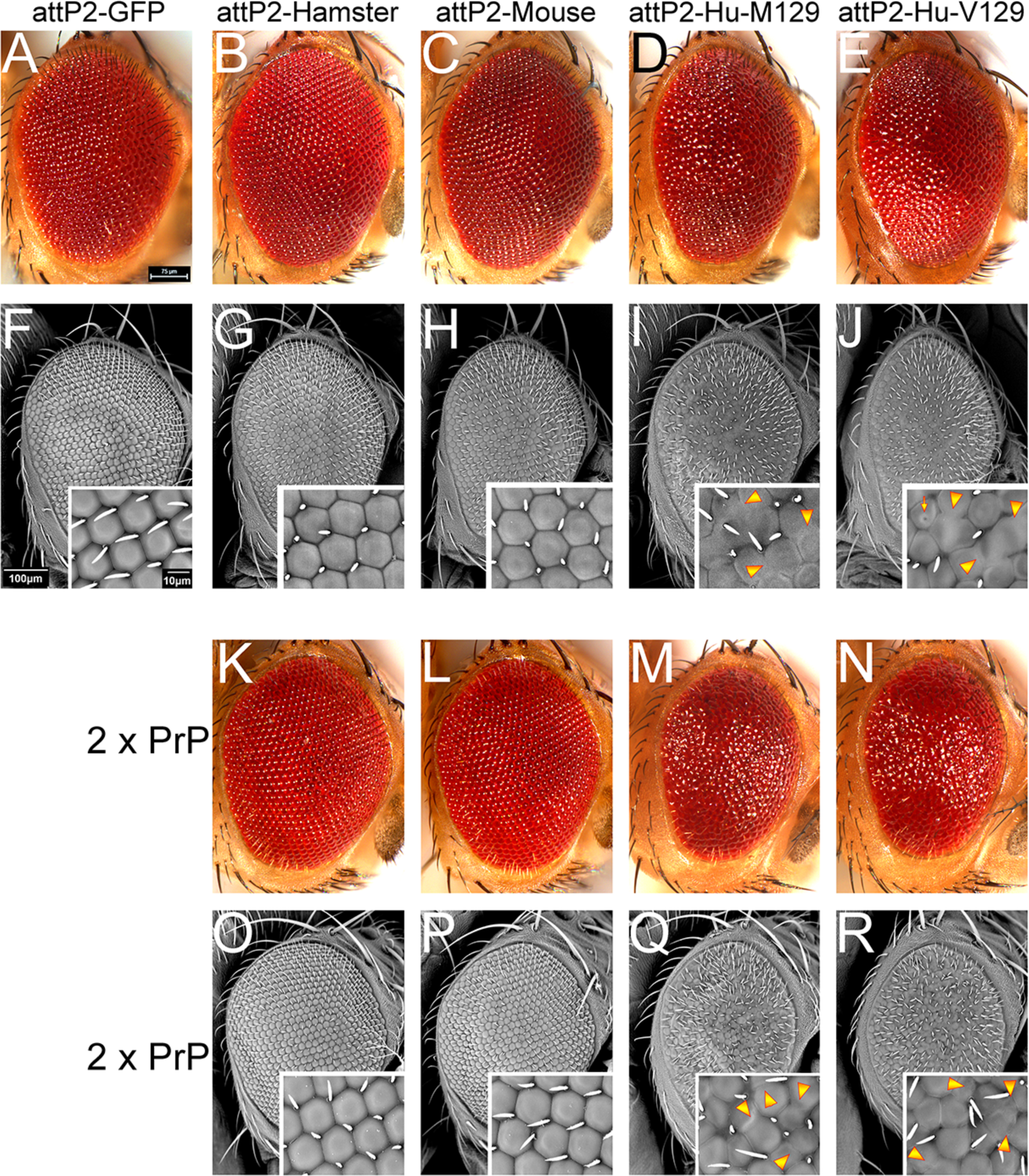
Human PrP-attP2 constructs exert heightened toxicity compared to rodent PrP-attP2. **A-E and K-N,** Fresh eyes and **F-J and O-R**, scanning electron micrographs of eyes expressing codon-optimized PrP constructs inserted in attP2. **A-J**, one copy of PrP constructs, **K-R**, two copies of PrP. **A and F**, Control eyes from flies expressing mCD8-GFP-attP2 under the control of *GMR-Gal4* (*GMR-Gal4 / UAS-mCD8- GFP-attP2*). Eyes are large and display a highly organized lattice. Flies expressing hamster (**B and G**) (*GMR-Gal4 / UAS-hamster PrP-attP2*) or mouse (**C and H**) (*GMR-Gal4 / UAS-mouse PrP-attP2*) PrP show normal eyes similar to those of control flies. **D and I**, Flies expressing human PrP-M129 (*GMR-Gal4 / UAS-human PrP-M129-attP2*) display slightly smaller and mildly disorganized eyes. **E and J**, Flies expressing human PrP-V129 (*GMR-Gal4 / UAS-human PrP-V129-attP2*) display smaller and highly disorganized eyes. **K-R**, Flies expressing two copies (2X) of the PrP constructs. Flies expressing two copies of hamster PrP (**K and O**), or mouse PrP (**L and P**), display large and organized eyes. **M and Q**, Flies expressing human PrP-M129 display slightly smaller and mildly disorganized eyes. **N and R**, Flies expressing human PrP-V129 display smaller and highly disorganized eyes.

Since the human attP2-PrP constructs induce mild eye phenotypes, it could be argued that rodent PrPs causes very mild eye phenotypes that could be detected by pushing their expression. To test this idea, we generated flies carrying two copies of the PrP-attP2 constructs with one copy of *GMR-Gal4*. Flies expressing two copies of the mouse or hamster PrP-attP2 still exhibit normal eyes (Fig 4K, L, O and P). High magnification micrographs show perfect arrangement of ommatidia (Fig. 4O and P, insets). In contrast, flies expressing two copies of human PrP-attP2 exhibit small and very disorganized eyes (Fig 4M, N, Q and R). The ommatidia are disorganized, have abnormal shapes, and appear highly fused (Fig. 4Q and R, insets). Thus, doubling the expression of rodent PrP still results in normal eyes, supporting not only the heightened toxicity of human PrP but the unique ability of human PrP to disrupt the fly eye.

### Expression analyses and subcellular distribution for the new <u>attP2</u> PrP lines

We examined the mRNA expression level for the new attP2-based lines by quantitative RT-PCR (qPCR). We generated homogenates from flies expressing attP2-PrP in the eye as described above, followed by mRNA extraction and examination of PrP expression by qPCR. The same primers were used for human PrP-M129 and -V129, but hamster and mouse PrP each had unique primers because of small sequence differences. All the primers were designed to amplify the same sequence. The internal control used for normalization was *Glyceraldehyde 3-phosphate dehydrogenase*. After normalization, all constructs showed identical expression levels (Fig. 5A), consistent with the shared landing site at attP2.

**Figure 5.**
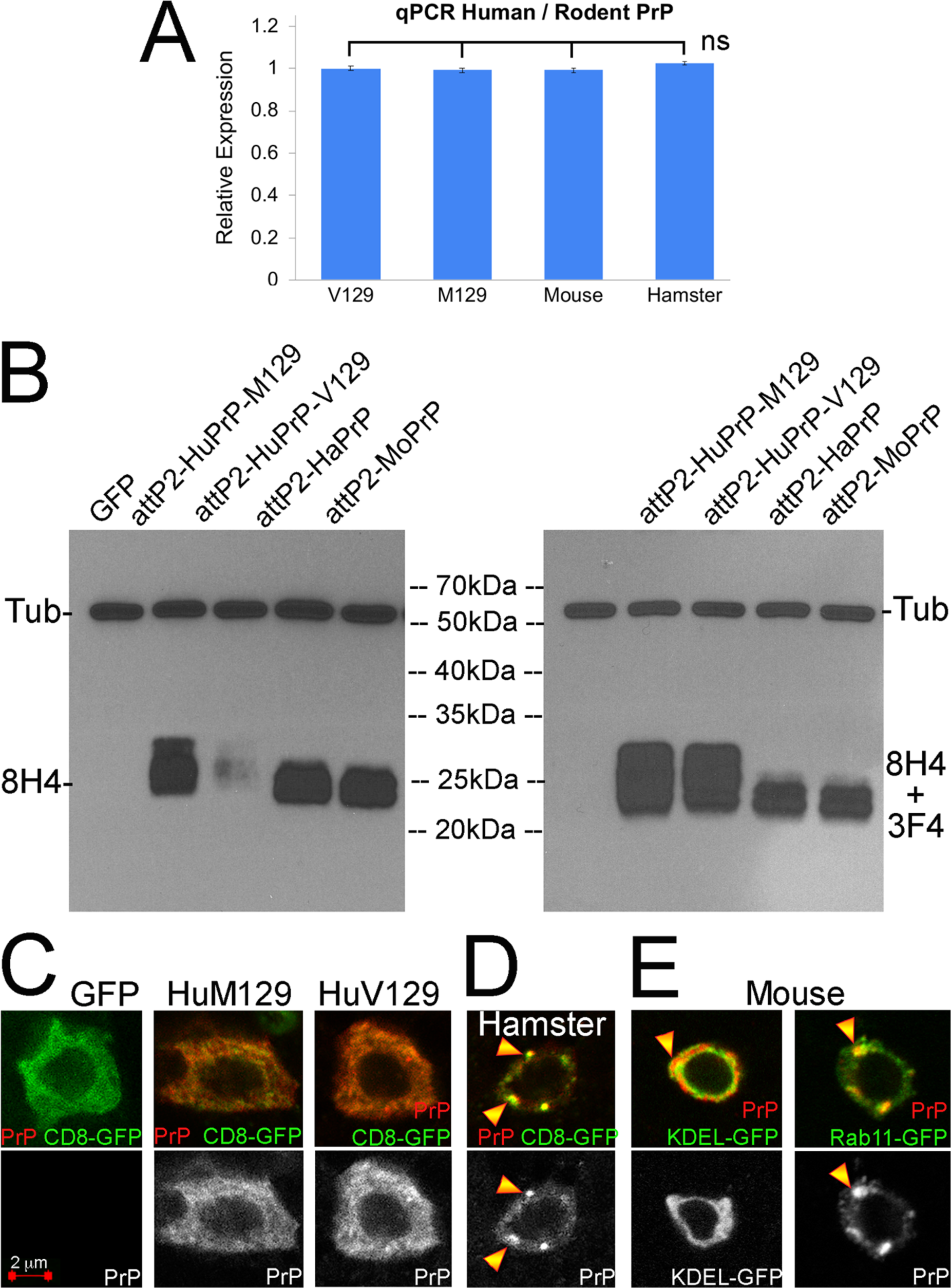
Human and rodent PrP undergo different biogenesis in flies. **A**, Identical expression levels of PrP mRNA in fly eyes by qPCR. **B**, PrP expression in fly eyes by western blot. Left, head-extracts from flies expressing GFP, human PrP-M129, human PrP-V129, mouse PrP, or hamster PrP in the eyes incubated with the 8H4 anti-PrP antibody and anti-Tubulin. Human PrP-M129 shows a distinct electrophoretic pattern compared to mouse PrP and hamster PrP but human PrP-V129-attP2 produces a weak signal. On the right, same membrane serially incubated with 8H4 and 3F4 antibodies showing normal levels of human PrP- V129. The pattern of human PrP-V129 is identical to that of human PrP-M129. **C-E**, Subcellular distribution of human PrP in *Drosophila* brain interneurons in the larvae ventral ganglion under the control of *OK107-Gal4*. C, Both human PrP-M129 and human PrP-V129 show a diffuse distribution through the ER that overlaps with mCD8-GFP. D and E, hamster and mouse PrP show punctate distribution that corresponds to post-ER secretory vesicles including Rab11 endosomes.

Next, we analyzed the new human and rodent PrP lines by western blot to identify differences at the protein level and the relative accumulation of isoforms. PrP has two facultative *N*-glycosylation sites and the relative usage of these two sites depends on their availability, which can be used to assess qualitative differences. We generated homogenates from flies expressing mCD8-GFP-attP2 or PrP-attP2 in the eye as described above. We first used the 8H4 anti-PrP antibody since this antibody binds to both human and rodent PrP. 8H4 revealed strong reactivity against human PrP-M129, hamster PrP, and mouse PrP, but showed a weak signal for human PrP-V129 (Fig. 5B, left panel). Note that all the lanes are equally loaded as indicated by the Tubulin signal present in the same gel at 50 kDa. This abnormal finding was consistent over multiple replicates. It is unlikely that V129 is expressed at very low levels compared to M129 since both induce similar eye phenotypes. Another possibility is that 8H4 detects a conformational difference between the M129 and V129 polymorphisms. Unfortunately, few antibodies detect conserved epitopes in human, hamster and mouse PrP, much less with the same affinity. We serially incubated the same membrane with 8H4 followed by 3F4, which recognizes human and hamster PrP, but not mouse PrP. The 8H4 incubation showed the same weak signal for human PrP-V129, but addition of 3F4 showed similar signal intensity and electrophoretic pattern for the two human PrP alleles (Fig. 5B, right panel). Both human PrPs present an additional band that runs higher than hamster and mouse PrP, suggesting that human PrP accumulates similar levels of the three glycoforms.

Last, we examined the subcellular distribution of PrP. We expressed all PrP-attP2 constructs under the control of *OK107-Gal4* and analyzed their expression in interneurons in the larval ventral ganglion. Both human PrP-M129 and -V129 show a diffuse distribution throughout the ER that colocalizes with membrane-bound CD8-GFP (Fig. 5C). Hamster and mouse PrP show a punctate distribution that partially colocalize with KDEL in the ER and Rab11 in recycling endosomes indicating abnormal maturation and secretion (Fig. 5D and E) (Fernandez-Funez et al., 2010, Fernandez-Funez et al., 2009). Overall, these differences in subcellular distribution and electrophoretic mobility indicate that human and rodent PrP undergo different posttranslational modifications and maturation through the secretory pathway. The accumulation of rodent PrP in the secretory pathway may contribute to its lower expression levels compared to human PrP and the weaker phenotypes described above.

### Extrinsic modifiers of PrP toxicity: interaction of human PrP and the amyloid-β peptide

We further tested the differences between human and rodent PrP by examining their genetic interactions. Multiple reports support the direct interaction of PrP and the amyloid-*β*42 (A*β*42) peptide in biochemical assays (Lauren et al., 2009, Chen et al., 2010, Zou et al., 2011, Gimbel et al., 2010, Gunther and Strittmatter, 2010, Balducci et al., 2010). PrP seems to be required for the manifestation of A*β* phenotypes in brain neurons in mouse models, suggesting a possible functional relationship between the molecular mechanisms mediating Alzheimer’s and prion diseases. The new PrP-attP2 lines allowed us to determine the functional interactions of human and rodent PrP with A*β*42. Since A*β*42 has robust eye phenotypes when expressed at high temperatures (27°C) (Casas-Tinto et al., 2011), we examined the potential interactions at 25°C to detect interactions resulting in enhanced eye phenotypes. As expected, flies expressing hamster and mouse PrP-attP2 under these mild conditions have normal eyes comparable to those expressing GFP (Fig. S1A-C). Expression of human PrP-M129-attP2 or -V129-attP2 results in subtle disruption of the ommatidia lattice (Fig. S1D and E). Co-expression of A*β*42 and GFP results in moderately disorganized eyes with a few black spots (Fig. S1F). Co-expression of hamster and mouse PrP with A*β*42 results in eyes similar to those of control flies (Fig. S1G and H). Remarkably, co-expression of human PrP- M129 or V129 with A*β*42 results in smaller and highly disorganized (glassy) eyes (Fig. S1I and J), demonstrating a specific functional interaction of human PrP and A*β*42 that is not observed between rodent PrP and A*β*42.

### Extrinsic modifiers of PrP toxicity: the unfolded protein response (UPR)

One of the best understood mechanisms mediating the toxicity of PrP is the accumulation of misfolded conformations in the ER, which cause ER stress and results in the activation of the UPR. The UPR encompasses the coordinated activity of three main pathways with homeostatic functions that can be activated during development and has a key protective role during acute cellular stress (Hetz et al., 2005, Moreno et al., 2012). The three ER membrane anchored sensors (PERK, Ire1*α*, and ATF6) are bound to Hsc3-70 / BiP under normal conditions in their inactive state. An increase in misfolded protein load in the ER results in detachment of BiP from the sensors, activating their downstream effectors. Ire1*α* activation results in the splicing of a small intron in the X-box binding protein 1 (XBP1) that causes a frame shift that activates XBP1s. We showed previously that expression of A*β*42 in flies activates Ire1*α* by using the XBP1- GFP sensor (Casas-Tinto et al., 2011, Ryoo et al., 2007). We used the same sensor here to examine the ability of human PrP to activate the Ire1*α* branch of the UPR. As control, we show here again that the UPR sensor XBP-GFP has very low basal activity and A*β*42 causes splicing and accumulation of GFP in the eye imaginal disc (Fig. 6A and B). Expression of random human PrP-V129 also induced UPR (Fig. 6C), but both mean levels of GFP and the integrated intensity of the fluorescent signal are significantly lower than those for A*β*42 (Fig. 6D). Combinations of human PrP with RNAi alleles for Ire1*α* and XBP1 resulted in small eyes despite these alleles having no effect on their own (Fig. 6E-G and J-L). Particularly striking is the combination with the XBP1 RNAi that results in a very small eye, indicating a key protective role of XBP1.

**Figure 6.**
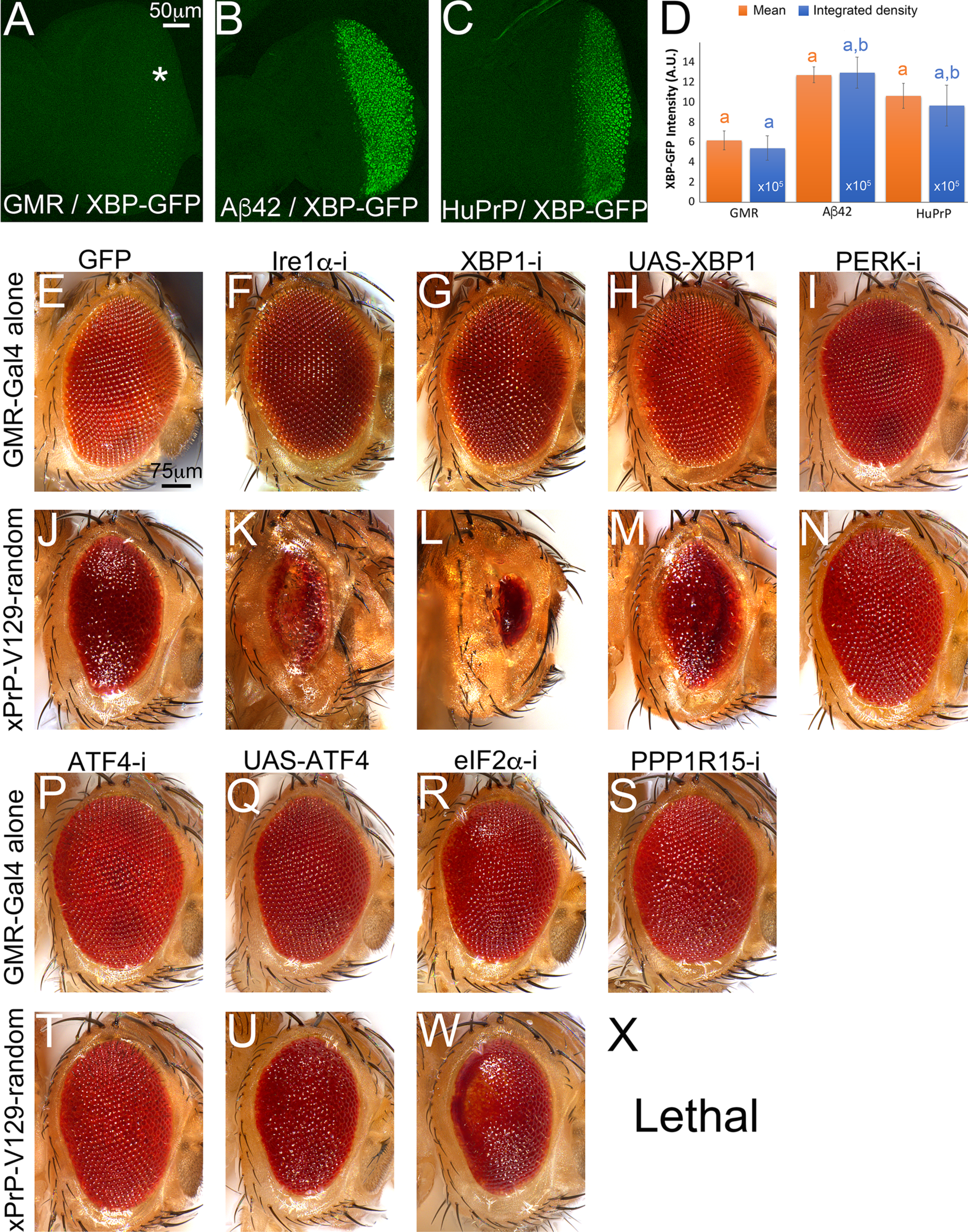
Silencing the PERK pathway suppresses human PrP toxicity. **A-D,** Human PrP activates the Ire1*α* branch of the UPR. Both human PrP and A*β*42 activate the XBP-GFP sensor, but A*β*42 is a stronger activator. Scale for the integrated density is x10^5^. All differences for mean intensity or integrated density are statistically significant by T-Test. a: *p* < 10^-5^; b: *p* < 10^-3^. **E-X**, Micrographs of fresh eyes from control flies expressing mCD8-GFP or UPR alleles alone (E-I and P-S) or in combination with human PrP-V129 (J-N and T-X) under the control of *GMR-Gal4* at 27°C. **E and J**, Control flies expressing mCD8-GFP alone (*GMR-Gal4 / UAS-mCD8-GFP-attP2*) or co-expressing human PrP-V129 (*GMR-Gal4 / UAS-human PrP- V129*) in the eye. **F and G,** Flies expressing Ire1*α*-RNAi (*GMR-Gal4 / UAS-Ire1α-RNAi*) or XBP1-RNAi (*GMR-Gal4 / UAS-XBP1-RNAi*) alone in the eye exhibit normal eyes. **K and L,** Flies co-expressing human PrP-V129 and Ire1*α*-RNAi (*GMR-Gal4 / UAS-Ire1α-RNAi / UAS-human PrP-V129*) or XBP1-RNAi (*GMR-Gal4 / UAS-XBP1-RNAi / UAS-human PrP-V129*) exhibit small and disorganized eyes. **H,** Flies expressing XBP1 alone (*GMR-Gal4 / UAS-XBP1*) have normal eyes. **U,** Flies co-expressing human PrP- V129 and XBP1 (*GMR-Gal4 / UAS-XBP1 / UAS-human PrP-V129*) show eyes comparable to controls. **I and P**, Flies expressing PERK-RNAi (*GMR-Gal4 / UAS-PEK-RNAi*) or ATF4-RNAi (*GMR-Gal4 / UAS- ATF4-RNAi*) alone in the eye exhibit normal eyes. **N and T,** Flies co-expressing human PrP-V129 and PERK-RNAi (*GMR-Gal4 / UAS-PEK-RNAi / UAS-human PrP-V129*) or ATF4-RNAi (*GMR-Gal4 / UAS- ATF4-RNAi / UAS-human PrP-V129*) exhibit large and well-organized eyes. **Q,** Flies expressing ATF4 alone (*GMR-Gal4 / UAS-ATF4*) have normal eyes. **U,** Flies co-expressing human PrP-V129 and ATF4 (*GMR-Gal4 / UAS-ATF4 / UAS-human PrP-V129*) show mildly suppressed eye organization. **R**, Flies expressing eIF2*α*-RNAi alone (*GMR-Gal4 / UAS-eIF2α-RNAi*) exhibit mildly disorganized eyes. **W,** Flies co-expressing human PrP-V129 and eIF2*α*-RNAi (*GMR-Gal4 / UAS-eIF2α-RNAi / UAS-human PrP- V129*) exhibit highly disorganized and mildly depigmented eyes. **S**, Flies expressing PPP1R15-RNAi alone (*GMR-Gal4 / UAS-PPP1R15-RNAi*) show mildly disorganized eyes. **X,** Flies co-expressing human PrP- V129 and PPP1R15-RNAi (*GMR-Gal4 / UAS-PPP1R15-RNAi / UAS-human PrP-V129*) results in pupal lethality.

The PERK branch is the most complex because it mediates both protective and maladaptive responses. Activated PERK phosphorylates eIF2*α* and prevents the interaction of the eIF2 complex with the ribosome, resulting in global translation inhibition. Acute phospho-eIF2*α* helps resolve ER stress by decreasing protein translation load; however, chronic ER stress can result in cell death. To resolve acute ER stress, unconventional translation of ATF4 (crc in flies) results in the transcriptional regulation of stress response genes and the PPP1R15 phosphatase (GADD34 in mammals). PPP1R15 dephosphorylates eIF2*α* to restore translation and prevent cell death from translation inhibition. In flies, PPP1R15 is directly regulated by phospho-eIF2*α*, but still plays the same feedback role to restore translation (Malzer et al., 2013). Recent reports highlighted the ability of PrP to chronically phosphorylate eIF2*α* and show the protective activity of genetic and pharmacological inhibition of PERK in prion-infected mice (Hughes and Mallucci, 2019, Moreno et al., 2012). We next examined the consequence of modulating the ability of PERK, ATF4 and to modulate the toxicity of human PrP. Silencing PERK or ATF4 alone has no effect in the eye (Fig. 6I and P). Remarkably, silencing PERK or ATF4 robustly suppressed PrP toxicity in the eye (Fig. 6N and T). We validated this result with multiple RNAi lines (*PEK^KK100348^, PEK^HMJ02063^, PEK^GL00030^*, *crc^KK111018^*, *crc^JF02007^*). In our hands, PERK overexpression is pupal lethal, although it has been shown to robustly disrupt eye development (Malzer et al., 2010). Overexpression of ATF4 alone results in normal eyes (Fig. 6Q) is mildly protective of PrP (Fig. 6U). Silencing eIF2*α* alone results in slight disorganization in the posterior eye (Fig. 6R) and enhanced the toxicity of PrP resulting in smaller, more disorganized, and partially depigmented eyes (Fig. 6W). Lastly, PPP1R15 silencing alone results in slightly disorganized eyes (Fig. 6S) but caused synthetic pupal lethality in combination with PrP with two different RNAi (*PPP1R15^KK104106^*, *PPP1R15^HMS00811^*) (Fig. 6X). This is consistent with a significant increase in the levels of phospho-eIF2*α* and inhibition of protein translation. These observations indicate that phospho-eIF2*α* is a main driver of PrP toxicity in flies. The suppression by ATF4-RNAi suggests that downstream effectors of ATF4 are also responsible for PrP toxicity. These results highlight the conserved molecular mechanisms of pathogenesis in mammalian and *Drosophila* models of PrP.

### Intrinsic mediators of toxicity: protective substitutions from animals resistant to prion diseases

As discussed above, several animals are recognized for their high natural resistance to prion diseases, including dogs, horses, rabbits, and pigs (Kirkwood and Cunningham, 1994, Espinosa et al., 2020, Vidal et al., 2020, Chianini et al., 2012, Bian et al., 2017). The amino acid sequence for PrP in these resistant animals have multiple differences compared to human PrP but it is unclear which mutations are protective and which are neutral (Fig. 7A). Structural studies identified amino acids proposed to mediate the structural stability resistant PrPs: D159 in dog, S167 in horse, and S174 in rabbit and pig (Myers et al., 2020, Khan et al., 2010, Perez et al., 2010, Lysek et al., 2005). These three residues are in a region encompassing the *α*1-*β*2 and *β*2-*α*2 loops (Fig. 7A). The 3D alignment of the globular domains of human, dog, horse, and rabbit PrP (Fig. 7B and C) shows high overall conservation. Some relevant differences include the shorter or lack of *β*-sheets in dog, rabbit and horse PrP compared to human PrP (Fig. 7B and C). Helix 2 is also longer in horse and rabbit PrP, and the CT3D domain exhibits clear differences among the four structures, although no clear structure-function correlation exists at this time (Fig. 7B and C). We hypothesize that these three residues impact the dynamics of the CT3D domain in their corresponding PrPs and are responsible for the different toxicity of human PrP compared with dog, rabbit, and horse PrP.

**Figure 7.**
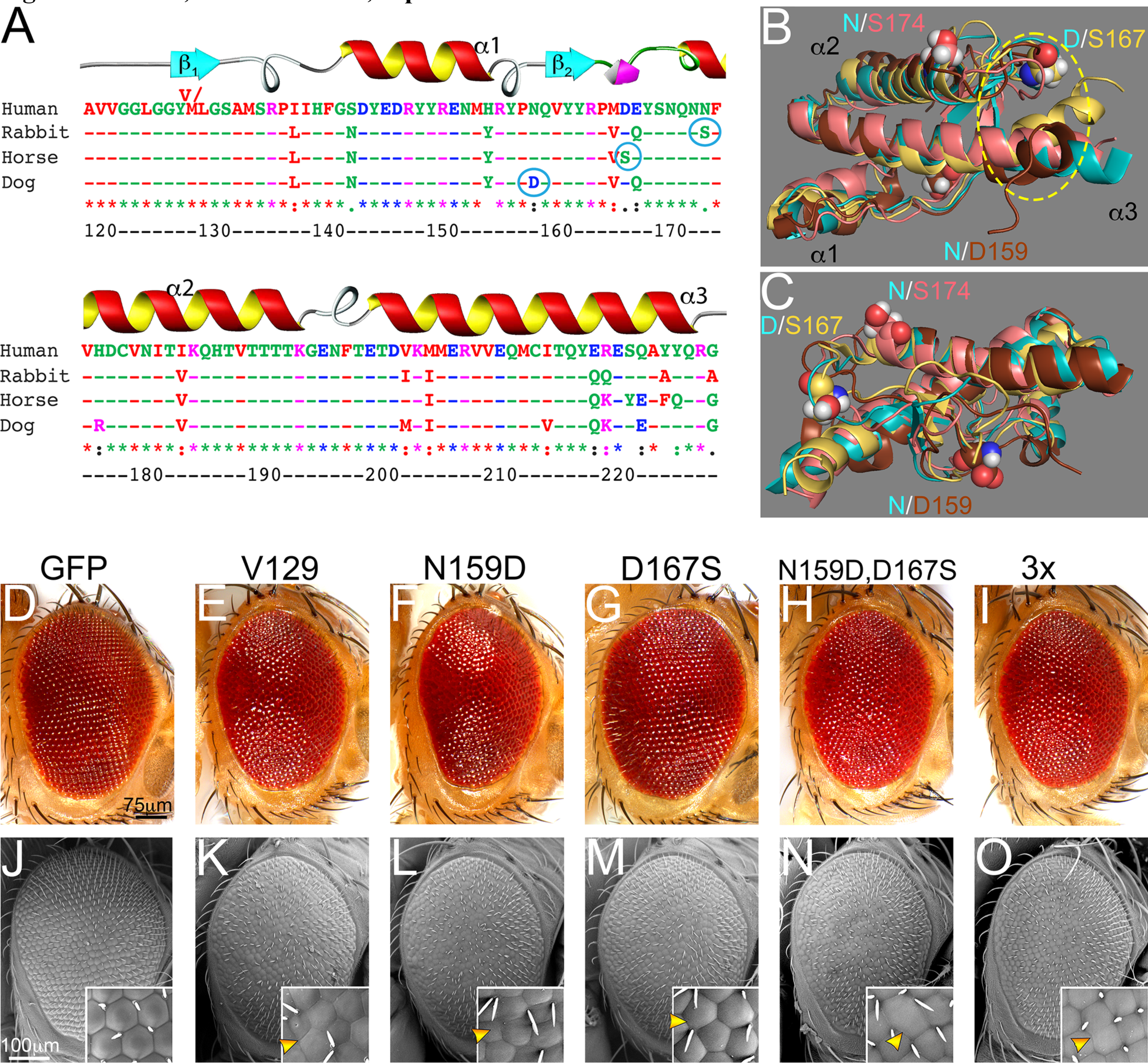
D167S is protective in the context of human PrP. **A**, **A,** Sequence alignment of the C-terminal globular domain of PrP from human, dog, horse, and rabbit. Amino acid numbering corresponds to human PrP. Candidate protective residues from rabbit, horse, and dog PrP are circled. **B and C**, 3D alignment of the globular domain of human (cyan), dog (brown), horse (yellow), and rabbit (salmon) PrP. The position of residues at 159, 167, and 174 are shown with the human amino acid in cyan and the corresponding color for each animal. **D-O**, Micrographs of fresh eyes (D-I) and scanning electron microscope (J-O) from control flies or flies expressing human PrP-attP2 constructs. **D and J**, Flies expressing a control reporter in the eye (*GMR-Gal4/UAS-mCD8-GFP-attP2*) show highly organized eyes. **E and K**, Flies expressing a human PrP- WT (V129 polymorphism) in the eye (*GMR-Gal4/UAS-human PrP-V129-attP2*) show disorganized, glassy eyes. Notice the poor differentiation and fusion of ommatidia in the inset (arrowhead). **F and L**, Flies expressing a human PrP-N159D in the eye (*GMR-Gal4/UAS-human PrP-N159D-attP2*) show disorganized and glassy eyes, with abnormal ommatidia (arrowhead). **G and M**, Flies expressing a human PrP-D167S in the eye (*GMR-Gal4/UAS-human PrP-D167S-attP2*) show partially rescued eye organization. Notice the improved arrangement of ommatidia (arrowhead). **H, I, N and O**, Flies expressing the 2X human PrP (*GMR-Gal4/UAS-human PrP-N159D,D167S-attP2*) or the 3x human PrP (*GMR-Gal4/UAS-human PrP- N159D,D167S,N174S-attP2*) in the eyes show partially rescued eye organization. Some abnormal ommatidia are still found (arrowheads).

### In vivo activity of protective substitutions: eye phenotype

We previously examined the consequence of introducing the equivalent amino acid substitution from human PrP into dog, horse, and rabbit PrP. Those experiments revealed that the single substitutions dog PrP-D159N and horse PrP-S167D become highly toxic in the *Drosophila* brain, whereas rabbit PrP-S174N had no effect (Sanchez-Garcia and Fernandez-Funez, 2018). To examine the mechanisms mediating human PrP toxicity, we next introduced the three protective residues from dog, horse, and rabbit PrP into human PrP-V129. We introduced N159D and D167S alone, combined (2x-N159D-D167S), or combined with N174S (3x-N159D-D167S-N174S). The N174S substitution alone is shown in a different manuscript together with Y225A (RMM and PFF, submitted). We generated transgenic flies by the same methods described above (codon-optimized and inserted in <u>attP2</u>) into the human PrP-V129 backbone.

Flies expressing human PrP-V129-attP2 in the eye at 27°C exhibit slightly smaller and moderately disorganized eyes compared to control flies expressing GFP-attP2 (Fig. 7D, E, J and K). Ommatidia appear disorganized and fused throughout the surface (Fig. 7K, inset). Flies expressing human PrP-N159D-attP2 show eyes similarly disorganized to those expressing V129 (Fig. 7F and L). High magnification shows the same differentiation problems as the flies expressing V129 (Fig. 7L, inset). Flies expressing human PrP- D167S-attP2 exhibit larger and better organized eyes than those expressing V129 (Fig. 7G and M). High magnification shows more definition of ommatidia, although they are abnormal (Fig. 7M, inset). Flies expressing the double mutant N159D-D167S and the triple mutant exhibit similar organization to the D167S mutant alone (Fig. 7H, I, N and O), indicating that the combinations showed no cooperative activity. Overall, these experiments show that N159D alone is not protective in the context of human PrP, whereas D167S is partially protective but shows no cooperativity with N159D and N174S.

### In vivo activity of protective substitutions: degeneration of brain neurons

We lastly examined the consequence of expressing the new attP2-based constructs in the mushroom bodies. Figure 8 shows the axonal projections of the mushroom body neurons at high magnification. We measured the surface of the projections in each genotype in young (day 1 post-eclosion) and old (day 40) flies to determine differences in size and shape. Control 1-day-old control flies show robust axonal projections, which split into dorsal (*α*) and medial (*β* and *γ*) lobes (Fig. 8A). Control 40-day-old flies preserve the normal shape but expand in surface (Fig. 8G, M and N), as we reported before (Sanchez-Garcia and Fernandez-Funez, 2018, Fernandez-Funez et al., 2010). 1-day-old flies expressing PrP-V129 exhibit thinner axonal projections with a significant difference with control flies (Fig. 8B and M). By day 40 these flies show extensive degeneration as indicated by the loss of *α* lobes and widespread membrane blebbing (Fig. 8H and M). 1-day-old flies expressing human PrP-N159D, D167S, 2X or 3X mutants exhibit similar axonal projections compared to young flies expressing PrP-V129 (Fig. 8C-F and M). All the mutants show extensive blebbing at day 40 but the preservation of the lobes is different. 40-day-old flies expressing N159D show similar area than controls expressing V129 (Fig. 8I, M and N), whereas older flies expressing D167S or 2X exhibit significantly larger lobes (Fig. 8J, M and N). Lastly, flies expressing the 3X mutant are the only showing an expansion of the mushroom body lobes as they age (Fig. 8L, M and N), but are still smaller than in controls. Overall, the analysis of the mushroom bodies shows that human PrP is highly toxic to brain neurons starting during development and continuing with extensive degeneration during aging, but constructs carrying the D167S substitution show moderate protection.

**Figure 8.**
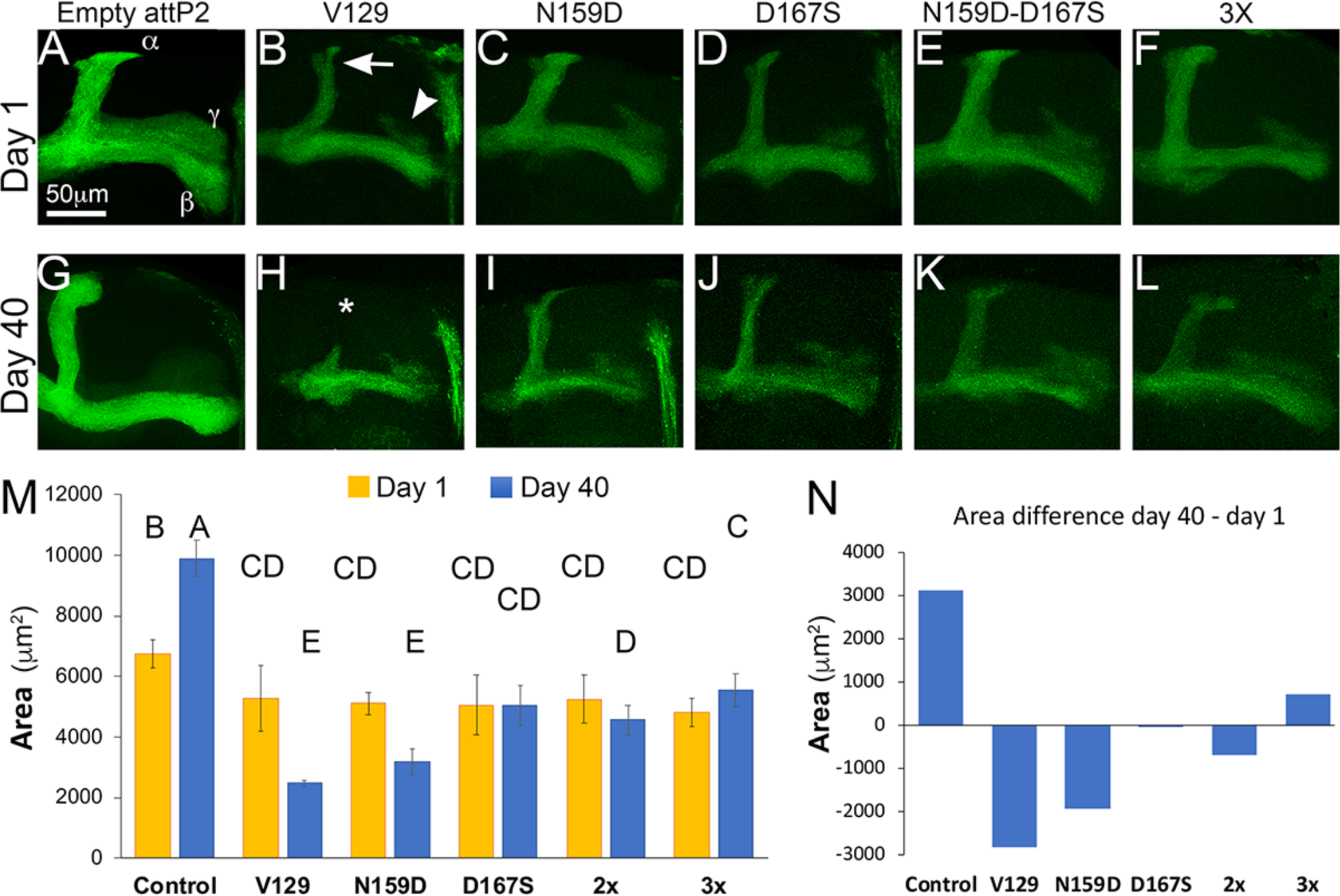
In vivo analysis of protective mutations in brain neurons. **A-L,** Micrographs of mushroom body axonal projections at days 1 (A-F) or 40 (G-L). **A and G**, 1- and 40-day-old flies carrying an empty attP2 site (*OK107-Gal4/UAS-mCD8-GFP/attP2*) show highly organized mushroom body axonal projections. The *α*, *β*, and *γ* lobes are indicated. 40-day-old flies show an increase in the surface of the projections. **B and H**, 1- and 40-day-old flies expressing a human PrP-WT (V129 polymorphism) (*OK107- Gal4/UAS-mCD8-GFP /UAS-human PrP-V129-attP2*) show thin projections at day 1 and significant degeneration by day 40, including the loss of the *α* lobe (H, *). **C and I**, 1- and 40-day-old flies expressing a human PrP-N159D (*OK107-Gal4/UAS-mCD8-GFP/UAS-human PrP-N159D-attP2*) show small projections at day 1 that continue to degenerate during aging. **D and J**, 1- and 40-day-old flies expressing a human PrP-D167S (*OK107-Gal4/UAS-mCD8-GFP/UAS-human PrP-D167S-attP2*) show small projections at day 1 but slower degeneration over time. **E, F, K and L**, 1- and 40-day-old flies expressing the 2X human PrP (*OK107-Gal4/UAS-mCD8-GFP/UAS-human PrP-N159D,D167S-attP2*) or the 3x human PrP (*OK107-Gal4/UAS-mCD8-GFP/UAS-human PrP-N159D,D167S,N174S-attP2*) show small projections at day 1 but slower degeneration over time. The 3X mutant is the only PrP that allows the projections to expand as the flies age. **M**, Quantification of axonal projections and statistical analysis. Statistical significance between groups is shown by the connecting letters. Levels not connected by the same letter are significantly different. P-value for different letter groups is <0.0001 except for B and C (p = 0.0037) and between CD and D (p = 0.021). **N**, Area differential day 40 – day 1 for each condition. Only the control flies and flies expressing the 3X mutant show an expansion of axonal projections over time.

## Discussion

Here we describe the characterization of new genetic tools to dissect the mechanism underlying PrP toxicity. All the new PrP constructs are codon-optimized for *Drosophila* and integrated in the same attP2 landing site. These optimized attP2 constructs are weaker than the random insertions but still induce an eye phenotype that critically supports the discovery of intrinsic and extrinsic mediators of PrP toxicity. Our findings suggest that human and rodent PrP acquire different conformations during biogenesis that affect maturation through the ER and Golgi, resulting in longer membrane expression and half-life for human PrP. This higher expression alone could explain the novel eye phenotype in flies expressing human PrP. Since doubling the dose of rodent PrP still results in normal eyes, we are confident that the novel phenotypes induced by human PrP are due to its intrinsic properties. In addition to the eye phenotype, random human PrP lines induced other specific phenotypes: vacuolation of photoreceptors, lethality following pan-neural expression, aggressive locomotor dysfunction under conditional expression, and complete elimination of the mushroom bodies in the adult brain. Importantly, human PrP expressed in flies is sensitive to mild PK digestion, indicating that no spontaneous accumulation of PrP^Sc^ occurs. Thus, PrP can be highly toxic without the PrP^Sc^ conformation, supporting the idea that neurotoxicity and transmission are caused by different PrP conformations (Sandberg et al., 2014, Sandberg et al., 2011). The lack of spontaneous PrP^Sc^ accumulation suggests that we can responsibly work with these flies at enhanced Animal Biosafety Level 2 (ABSL2) instead of ABSL3.

As a proof-of-concept for the extrinsic factors regulating PrP toxicity, we examined the functional interaction of human PrP with A*β*42 and the UPR. In 2009, A*β*42 was found to bind the unstructured N-terminal domain of PrP, a novel interaction proposed to mediate A*β*42-dependent inhibition of long-term potentiation (Lauren et al., 2009). Despite some initial resistance from the prion community (Calella et al., 2010, Kessels et al., 2010, Balducci et al., 2010), the interaction was confirmed by different techniques, although some studies still disagree on whether this interaction mediates A*β*42 toxicity (Chen et al., 2010, Zou et al., 2011, Gimbel et al., 2010, Gunther and Strittmatter, 2010, Balducci et al., 2010). The native PrP conformation has been proposed to work as a scaffold in lipid rafts that brings together several membrane proteins, including glutamate and lamin receptors (Zhang et al., 2019). The A*β*42 – PrP interaction stimulates glutamate receptors whereas the interaction with lamin receptors internalizes the complexes, providing access to the ER by retrograde transport, where A*β*42 causes significant ER stress (Casas-Tinto et al., 2011). Here, we show that A*β*42 and human PrP, but not hamster or mouse PrP, combine to increase their toxicity. This specific interaction could explain the contradictory results in the literature. We previously showed that human PrP has more binding sites for A*β*42 (six) than mouse PrP (one), supporting the functional interaction observed here (Zou et al., 2011). Additionally, A*β*42 and human PrP induce similar, although not identical, eye phenotypes in flies suggesting that A*β*42 and PrP perturb similar gene networks in the eye, including ER stress.

Our candidate extrinsic modifiers showed that silencing either PERK or its effector ATF4 robustly suppress the toxicity of human PrP, whereas silencing Ire1*α* or XBP1 robustly enhance it. Human PrP induces the Ire1*α* or XBP1 branch of the UPR in flies and the activity of this pathway is clearly protective since reduced Ire1*α* or XBP1 function enhances PrP toxicity. It is surprising, though, that overexpression of XBP1 has no effect on PrP toxicity,. This could be explained by interactions with the PERK pathway, which shuts down translation and limits the transcriptional response of XBP1. The robust protective activity of lowering PERK is consistent with recent findings in prion infected mice (Hughes and Mallucci, 2019, Moreno et al., 2012). These results suggest that phospho-eIF2*α* is a major mediator of PrP toxicity through its ability to block translation. In this line, we found that lowering the levels of PPP1R15, which leaves eIF2*α* in the phosphorylated state, caused synthetic lethality in combination with PrP. In contrast, PPP1R15 overexpression alone is not toxic in the absence of PrP, suggesting that normal eye development has low levels of phospho-eIF2*α*. A couple of our observations do not directly fit into the model in which PrP induces toxicity mostly through phospho-eIF2*α*. First, silencing eIF2*α* shows additive eye phenotypes in combination with PrP rather than causing lethality, suggesting that other downstream effectors mediate the toxicity. More importantly, the robust suppression of PrP toxicity by ATF4 silencing indicates that effectors downstream of ATF4 also participate in the maladaptive response. An important difference in the UPR is that in flies PPP1R15 is not a transcriptional target of ATF4 and, therefore, has no ability to directly modulate eIF2*α* phosphorylation. Hence, factors downstream of ATF4 could mediate the maladaptive response of the PERK branch. Recent studies have identified another translational regulator, 4E-binding protein (4E-BP), as an ATF4 transcriptional target that binds eIF4E and prevents the assembly of the eIF4F complex, which is critical for the entry of capped mRNAs into the ribosomal small subunit. Thus, a second mechanism converging into ribosomal activity may mediate the protective effects of silencing PERK and ATF4, and idea that we will investigate in follow up studies.

At this time, we do not yet fully understand the exact intrinsic mechanisms mediating the conformational dynamics of PrP and how they translate into different toxicity, disease susceptibility, or strain variability. While it seems clear that a few amino acid differences between mammalian PrPs are responsible for conformational differences, it remains challenging to pinpoint how specific amino acids contribute to PrP conformation (Myers et al., 2020). The new *Drosophila* model expressing human PrP enables mechanistic studies into sequence-structure-phenotype analyses through the efficient introduction of candidate mutations into the human PrP backbone. In a previous report we showed that two humanized mutants, dog PrP-D159N and horse PrP-S167D, turned toxic these non-toxic PrPs (Sanchez-Garcia and Fernandez-Funez, 2018). The D159N and S167D substitutions are proposed to increase conformational instability of the globular domain, misfolding and toxicity. Our prediction was that introducing the corresponding protective residues into human PrP would be protective, i.e., would revert the eye and mushroom bodies phenotypes. However, D167S is mildly protective whereas N159D has a weak effect in the mushroom bodies. Interestingly, the combinations N159D / D167S or the 3X mutant showed similar activity to D167S alone. These results provide valuable lessons about the rules governing PrP misfolding and toxicity. First, single amino acid changes are not enough to alter the high structural dynamics and toxicity of human PrP. In contrast, the corresponding amino acid changes transformed dog and horse into toxic isoforms. Second, N159D and D167S are not known to form distinct secondary or tertiary structures in dog and horse PrP (Perez et al., 2010, Lysek et al., 2005), suggesting that each alone will not introduce significant changes in conformational dynamics. S174 participates in a helix-capping domain that stabilizes helix 2 in rabbit PrP (Khan et al., 2010). However, addition of N174N in the 3X mutant had only a small effect observed in brain neurons. Third, combining amino acid changes from different animals did not increase the conformational stability of human PrP. A more likely strategy would consist of combining additional amino acid changes, including conservative changes *<u>from the same animal</u>* to recreate local structural features of dog, horse, or rabbit PrP. We are currently testing several such combinations, including Y225A from rabbit (PFF, submitted). The ability to efficiently test candidate mutations *in vivo* will eventually provide answers to the questions posed above about the genotype - morphotype - phenotype correlations.

## Materials and Methods

### Sequence alignment and 3D protein visualization

The alignments of the globular domain of human, hamster, mouse, dog, horse, and rabbit prion protein sequences was done using ClustalW2 (www.ebi.ac.uk/Tools/clustalw2). We used human PrP as reference and amino acid numbering for all species refers to the corresponding amino acid in human PrP (see Fig. 1A). Amino acid sequences were obtained from NCBI with the following accession numbers: AAH22532 (human), B34759 (Syrian hamster), and AAA39996 (mouse), AAD01554 (rabbit), ACG59277 (horse), and ACO71291 (dog). The color-coded amino acids indicate properties relevant for protein structure (size and charge). To generate 3D views of human, mouse and Syrian hamster PrP, we opened in PyMOL (pymol.org) the published NMR structures for human (1QM2), mouse (1XYX), hamster (1B10), rabbit (2FJ3), horse (2KU4), and dog (1XYK) PrP deposited in the RSCB Protein Data Bank (rcsb.org/pdb). We displayed the proteins in Cartoon formats showing only relevant amino acids to optimize their visualization. We also displayed the *β*2-*α*2 loop using the Surface and Mesh views.

### Generation of transgenic flies and genetics

#### Random insertions

Flies carrying the human PrP-WT (V129) construct in a random insertion were described previously (Fernandez-Funez et al., 2017). We generated flies carrying human PrP-M129 in a random insertion following the same procedures described above. <u>attP2 insertions</u>: The constructs carrying human PrP-M129, human PrP-V129, hamster PrP-WT, and mouse PrP-WT, and the human PrP mutants human PrP-N159D, -D167S, -N159D-D167S (double), and -N159D-D167S-N174S (triple, 3X) (all in V129 background) were chemically synthesized by Integrated DNA Technologies (IDT) using codon-optimized sequences for *Drosophila*. Assembled sequences were cloned between *XhoI* and *Xba*I sites onto the pJFRC7-20XUAS-IVS-mCD8:GFP *Drosophila* expression vector (Addgene #26220, (Pfeiffer et al., 2010)) after removing the mCD8:GFP transgene. The final constructs were sequenced to verify their integrity. The pUAST-based constructs were injected into *yw* embryos at Rainbow Transgenics following standard procedures (Rubin and Spradling, 1982) to generate multiple independent transgenic lines for each plasmid. Two independent strains were generated for each construct since they are all inserted in the same attP locus.

The driver strains *GMR-Gal4* (retina, all eye cells) (Mathew Freeman, Univ. of Oxford), *OK107-Gal4* (mushroom bodies) (Connolly et al., 1996), *Elav-Gal4* (pan-neural) (Lin and Goodman, 1994), *Elav-GS* (pan-neural, GeneSwitch) (Roman et al., 2001), the reporters *UAS-LacZ* and *UAS-mCD8-GFP*; the overexpression lines *UAS-PERK* (*pek*); and the TRiP RNAi lines *PEK^HMJ02063^, PEK^GL00030^*, *crc/ATF4^JF02007^*, and *PPP1R15^HMS00811^* were obtained from the Bloomington Drosophila Stock Center (<u>fly.bio.indiana.edu</u>). RNAi alleles for *PERK (v110278)*, *ATF4 (109014)*, *PPP1R15 (v107545)*, and *eIF2α (v104562)* were obtained from the Vienna Drosophila Stock Center (<u>stockcenter.vdrc.at/control/main</u>). Transgenic flies expressing human A*β*42 were described previously (Casas-Tinto et al., 2011) and the XBP-GFP sensor was obtained from HD Ryoo (Ryoo et al., 2007). UAS alleles for ATF4 was obtained from FlyORF (<u>flyorf.ch/index.php</u>). Fly stocks were maintained on standard *Drosophila* medium at 25°C. For experiments, homozygous females for the *Gal4* strains were crossed with *UAS* males to generate progeny expressing *PrP* in the desired tissue. Crosses were placed at 25°C for two days, transferred to 27°C until the progeny completed development, and adults were aged at 27°C, unless otherwise indicated.

### Characterization of eyes

We expressed all the constructs in the eye under the control of *GMR-Gal4*. Crosses were performed at 25°C for 2 d and the progeny were raised at 28°C, and adult flies were collected at day 1. To image fresh eyes, we froze the flies at -20°C for at least 24 h and collected images as z-stacks with a Leica Z16 APO using a 2X Plan-Apo objective. Flattened in-focus images were produced with the Montage Multifocus module of the Leica Application Software. For scanning electron microscopy, flies were serially dehydrated in ethanol, critically dried, and metal-coated for observation in a Jeol JSM-6490LV. For transmission electron microcopy, we collected flies of the appropriate genotype 1-day post eclosion, fixed the heads in 3% glutaraldehyde overnight, washed in phosphate buffer, post-fixed in 1% OsO4, dehydrated in ethanol and propylene oxide, embedded in resin, and subsequently mounted the heads in molds as described previously (Fernandez-Funez et al., 2000). Blocks were then cut into semithin sections (1 μm), stained with toluidine blue and imaged in a Nikon Eclipse Ni microscope with a 100x Plan Apo oil 1.4 NA objective. For ultrastructural analysis of the eyes, we collected ultrathin sections (70 nm), stained the sections, and imaged the samples between 2,500 and 25,000x magnifications using a Jeal JEM-1400PLUS TEM at the University Imaging Centers.

### Drosophila homogenates and western blot

Ten flies per genotype and time point were used for analysis. Fly heads were homogenized in 30 µl of RIPA buffer containing Complete protease inhibitors (Roche) using a motorized pestle and centrifuged for 1 min at 1,000 rpm. 25 µl of supernatant was mixed with loading buffer and resolved by SDS-PAGE in 4- 12% Bis–Tris gels (Invitrogen) under reducing conditions and electro-blotted onto nitrocellulose membranes. Membranes were blocked in TBS-T containing 5% non-fat milk and probed against the primary antibodies: anti-PrP 8H4 (1: 10,000, Millipore), anti-PrP 3F4 (1: 10,000, Prionics), anti-*β*-Tubulin (1: 50,000, Invitrogen). The secondary antibody was anti-Mouse-HRP (1: 4,000) (Jackson ImmunoResearch). Immunoreactive bands were visualized by enhanced chemiluminescence (ProSignal Dura ECL, Genesee). The protein biochemistry protocols are described in more detail in (Sanchez-Garcia et al., 2013). For the protease-resistance assay, fly brain homogenates were incubated with PK concentrations from 0 to 15 μg/ml for 30 min at 25°C. The digestions were stopped by adding 2mM PMSF and analyzed by western blot by staining with the 3F4 antibody.

### Quantitative RT-PCR (qPCR)

Ten male flies 1-2 days post eclosion were used per genotype for analysis. Fly heads were homogenized in 100 µl RTL buffer from RNeasy kit (Qiagen) using a motorized pestle. An additional 250 µl RTL were added and then centrifuged for 3 minutes at 14,000 rpm. Supernatant was collected, placed in a new tube, and used for RNA extraction using the RNeasy kit. Additional DNase (DNase I, NEB) treatment and ethanol precipitation was performed. Omniscript Reverse Transcription Kit (Qiagen) was used for cDNA synthesis, following the manufacturers protocol and using 50 ng RNA for each sample. cDNA was then diluted 5x before qPCR.

qPCR was performed on a Roche Lightcycler 480 Instrument II and using SYBR Green I Master Mix (Roche), following the manufacturers protocol. The housekeeping *Drosophila* gene *Glyceraldehyde 3- phosphate dehydrogenase* (*GAPDH*) was used as an internal control. Negative RT controls were run to eliminate contaminating genomic DNA. The following primers were used: Human PrP forward- GCGGCAATCGTTACCCTCCTC; Human PrP reverse- ACTGGGCTTATTCCACTGGGAGT; mouse PrP forward- GTAACCGCTACCCACCGCAAG; mouse PrP reverse- TGGTTTGCTGGGCTTGTTCCA; hamster PrP forward- TCCCCAGGAGGTAATCGGTATCCT; hamster PrP reverse- TGGTTATGAGTGCCTCCACCCT; GAPDH forward- TAAATTCGACTCGACTCACGGT; GAPDH reverse- CTCCACCACATACTCGGCTC. Each genotype had three biological replicates along with three technical replicates. The -ΔΔct method was used for data analysis and represented as the relative expression to human PrP.

### Immunofluorescence and microscopy

For subcellular localization, we co-expressed the *PrP* constructs and *mCD8-GFP* as a membrane marker in interneurons of the larval ventral ganglion under the control of *OK107-Gal4* (*UAS-mCD8-GFP; OK107-Gal4/UAS-PrP*). Whole-mount immunohistochemistry of fixed larval brains was conducted by fixing in 4% formaldehyde, washing with PBT, and blocking with 3% bovine serum albumin before incubating with the primary antibody as described previously (Fernandez-Funez et al., 2010). We incubated first with the 6H4 anti-PrP antibody (1: 1,000 dilution) followed by the secondary antibody anti-mouse- Cy3 (Molecular Probes) at 1:600 dilution. We mounted the stained larval brain in Vectashield antifade (Vector) mounting medium for microscopic observation and documentation. Images of whole brains were taken at 100X and single neurons were imaged at 630x (see details below). For mushroom body imaging, we crossed *OK107-Gal4; mCD8-GFP* flies with *LacZ* alone (negative control) or with *PrP* constructs (*UAS-mCD8-GFP; OK107-Gal4/UAS-PrP*) at 27°C. Adult flies were collected at days 1 and 40 post eclosion. We then imaged whole-mount adults brains labeled with mCD8-GFP at low magnification by dissecting, fixing, and mounting as described above. We imaged the entire brains as Z-stacks with the 10x objective and visualized the brain by flattening the Z-stacks (maximum intensity projections).

We collected fluorescent images in an LSM 710 Zeiss confocal system using 10X NA: 0.45 (air), 20X NA: 1.0 (air), and 63X NA: 1.4 (oil) objectives. We collected thick samples as Z-stacks and all genotypes were imaged using the same settings. From the Z-stacks, we created maximum intensity projections \ using the Zeiss Zen software. These images were combined using Adobe Photoshop; processing included trimming of non-informative edges and brightness / contrast adjustment to whole images. Signal intensity for XBP-GFP was extracted in Photoshop. The surface for mushroom body axonal projections was manually outlined in Adobe Photoshop 2021. Data was exported to Excel to calculate averages, standard deviations and created the graph. Oneway ANOVA analysis for mushroom bodies was conducted in JMP Pro 14. Following the finding that the averages were statistically significant, we performed a Tukey-Kramer post hoc pair-wise analysis of significance to determine which pairs were statistically different while reducing the false positive due to the analysis of multiple pairs. For XBP-GFP, we used pair-wise T-Test to determine differences between 3 samples. *p* < 0.05 was considered statistically significant.

### Behavior, locomotor assays

For the strong random human PrP insertion, we performed locomotor assays following conditional expression in adult flies with the Elav-GS system. For this, we combined *Elav-GS* with *UAS-LacZ*, *UAS- hamster PrP-random*, and *UAS-human PrP-random* and placed the crosses in fly media without the activator RU486 (Sigma). When the adult flies eclosed, we collected 20 females per replicate and split them in two groups: one in vials without RU486 and one with RU486. Then, we examined the ability to move vertically in an empty vial (climbing assay) at 28°C (Le Bourg and Lints, 1992). Briefly, 20 newborn adult females were placed in empty vials in duplicate and forced to the bottom by firmly tapping against the surface. After 10 sec, the number of flies that climb above 5 cm was recorded. This was repeated 10 times to obtain the average climbing index each day. At the end of the assay, the climbing index (flies above line/total flies x 100) was plotted as a function of age in Excel. Finally, data series for locomotor activity were analyzed for statistical significance by a 2-way ANOVA using JMP Pro 14. Since ANOVA showed strong significance for genotype and age, we calculated pair-wise *t*-test significance with Bonferroni multiple comparisons post-hoc test to minimize type I error. To avoid inflating differences between individual climbing sets, we first averaged the replicates and then averaged the climbing index for two consecutive days. Then, we performed pair-wise *t*-tests comparing each genotype to the control flies (LacZ) and each PrP mutant to its corresponding WT allele.

## Acknowledgements

We thank the Bloomington Drosophila Stock Center (NIH P40OD018537), the Vienna Drosophila Stock Center, and FlyORF, HD Ryoo for transgenic flies; the RSCB Protein Data Bank and ClustalW2, for free data and software; and the University of Minnesota Information Technology Support Services for institutional copies of PyMOL, Adobe products and JMP Pro 14. This work was supported by the resources and staff at the University of Minnesota Imaging Centers (SCR_020997). Gail Celio helped with sample preparation and TEM imaging of the eyes. This work was supported by the NIH grant 7R21NS096627-02 and the Winston and Maxine Wallin Neuroscience Discovery Fund award CON000000083928 to PF-F. Confocal and SEM images were collected at the Research Instrumentation Laboratory (UMN-UMD).

## Author contributions

PFF conceived the study; RM, JSG, DL and PFF performed experiments; RM, JSG, DL and PFF interpreted the experiments; RM, DL and PFF wrote the manuscript. All authors revised and approved the final manuscript.

---

**Figure S1.**
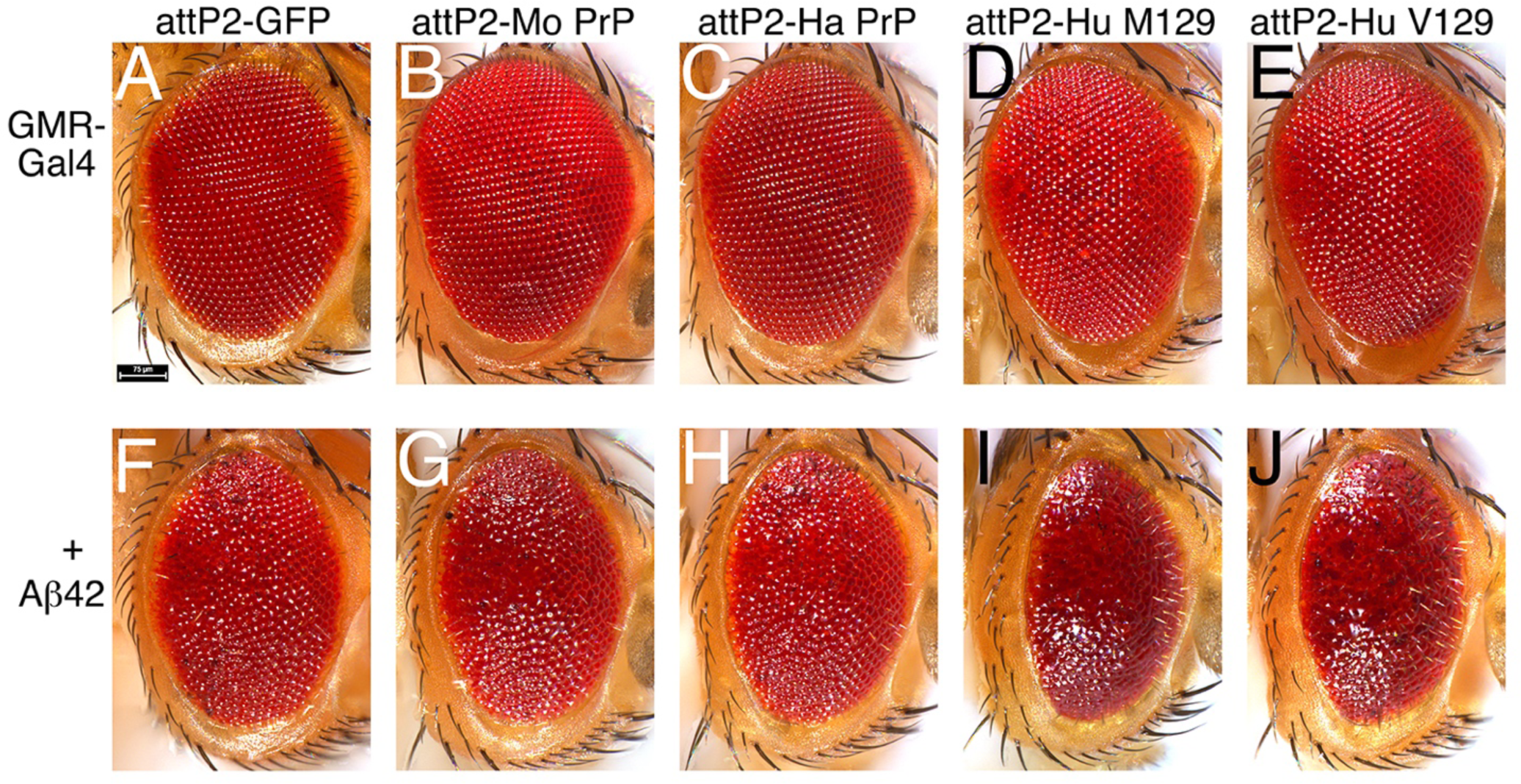
Human PrP enhances the toxicity of Aβ42. **A-J**, Micrographs of fresh eyes expressing mCD8- GFP, hamster PrP, mouse PrP, human PrP-M129, or human PrP-V129 alone (A-E) or in combination with A*β*42 (F-J) in the eye under the control of *GMR-Gal4* at 25°C. **A**, Control eyes from flies expressing mCD8- GFP (*GMR-Gal4 / UAS-mCD8-GFP-attP2*). **B and C**, Eyes from flies expressing mouse or hamster PrP (*GMR-Gal4 / UAS-mouse PrP-attP2 and GMR-Gal4 / UAS-hamster PrP-attP2*) are normal. **D and E**, Eyes from flies expressing human PrP (*GMR-Gal4 / UAS-human PrP-M129-attP2 and GMR-Gal4 / UAS-human PrP-V129-attP2*) show mild disorganization. These phenotypes are weak because the expression of PrP constructs is lower at 25°C. **F**, The eyes from flies co-expressing GFP and A*β*42 (*GMR-Gal4 / UAS-mCD8- GFP-attP2/ UAS-Aβ42*) are disorganized and have necrotic spots. **G and H**, The eyes from flies co-expressing rodent PrP with A*β*42 (*GMR-Gal4 / UAS-mouse PrP-attP2 / UAS-Aβ42 and GMR-Gal4 / UAS- hamster PrP-attP2 / UAS-Aβ42*) are similar to those in F. **I and J**, The eyes from flies co-expressing human PrP and A*β*42 (*GMR-Gal4 / UAS-human PrP-M129-attP2 / UAS-Aβ42 and GMR-Gal4 / UAS-human PrP- V129-attP2 / UAS-Aβ42*) are smaller and highly disorganized (glassy).

